# Basal cytoplasmic calcium levels regulate *C. elegans* germ stem cell proliferation

**DOI:** 10.1101/2025.10.03.680197

**Authors:** Alexandra C. Wells, Parva M. Vyas, Aahana N. Shankaran, Corrissa J. Velder, Rabia N. Kaya, Edward T. Kipreos

## Abstract

The *C. elegans* germline is one of the foremost model systems for the study of germ cell biology. We have created an integrated ratiometric calcium reporter that allows comparative analysis of calcium levels in germ cells. We observe a positive correlation between the rate of cell proliferation and calcium levels. Conditions that decrease proliferation decrease basal cytoplasmic calcium levels, these include starvation and loss of insulin, TGF-β, and folate signaling. Conversely, overproliferating germline tumor cells have increased basal cytoplasmic calcium levels. To determine if basal calcium levels regulate the rate of proliferation, we decreased calcium levels by inactivating the TRPM7-related GON-2 calcium channel. A *gon-2* null mutant has significantly decreased basal cytoplasmic calcium levels and germ cell proliferation. Partial reduction of GON-2 activity in a *gon-2* mutant heterozygote also reduces calcium levels and proliferation, but to a lower extent. This identifies GON-2 as a calcium channel that is required to maintain basal cytoplasmic calcium levels in germ cells. Conversely, increasing basal cytoplasmic calcium levels in germ cells either by inhibiting the plasma membrane calcium ATPase (which transports cytoplasmic calcium out of cells) or by inhibiting the SERCA channel (which transports cytoplasmic calcium into the endoplasmic reticulum) increased basal calcium levels and germ cell proliferation. This study identifies the germ line as the first tissue in *C. elegans* whose rate of cell proliferation is regulated by calcium. While transient increases in calcium are associated with cell proliferation in many animal species, our study suggests that the basal level of cytoplasmic calcium acts as a rheostat to modulate the rate of *C. elegans* germ cell proliferation in response to physiological processes and signals.

## Introduction

The *C. elegans* germ line is a model system for studying germ cell biology.^1^ The hermaphrodite germ line has two U-shaped arms, and contains both male and female germ cells and associated somatic gonad structures (Fig. 1A). The hermaphrodite gonad switches from male (sperm producing) to female (oocyte producing) states during the L4 larval and young adult stages. Sperm are stored in the spermatheca, a somatic gonadal structure between the uterus and the oviduct in the proximal gonad.

**Figure 1.**
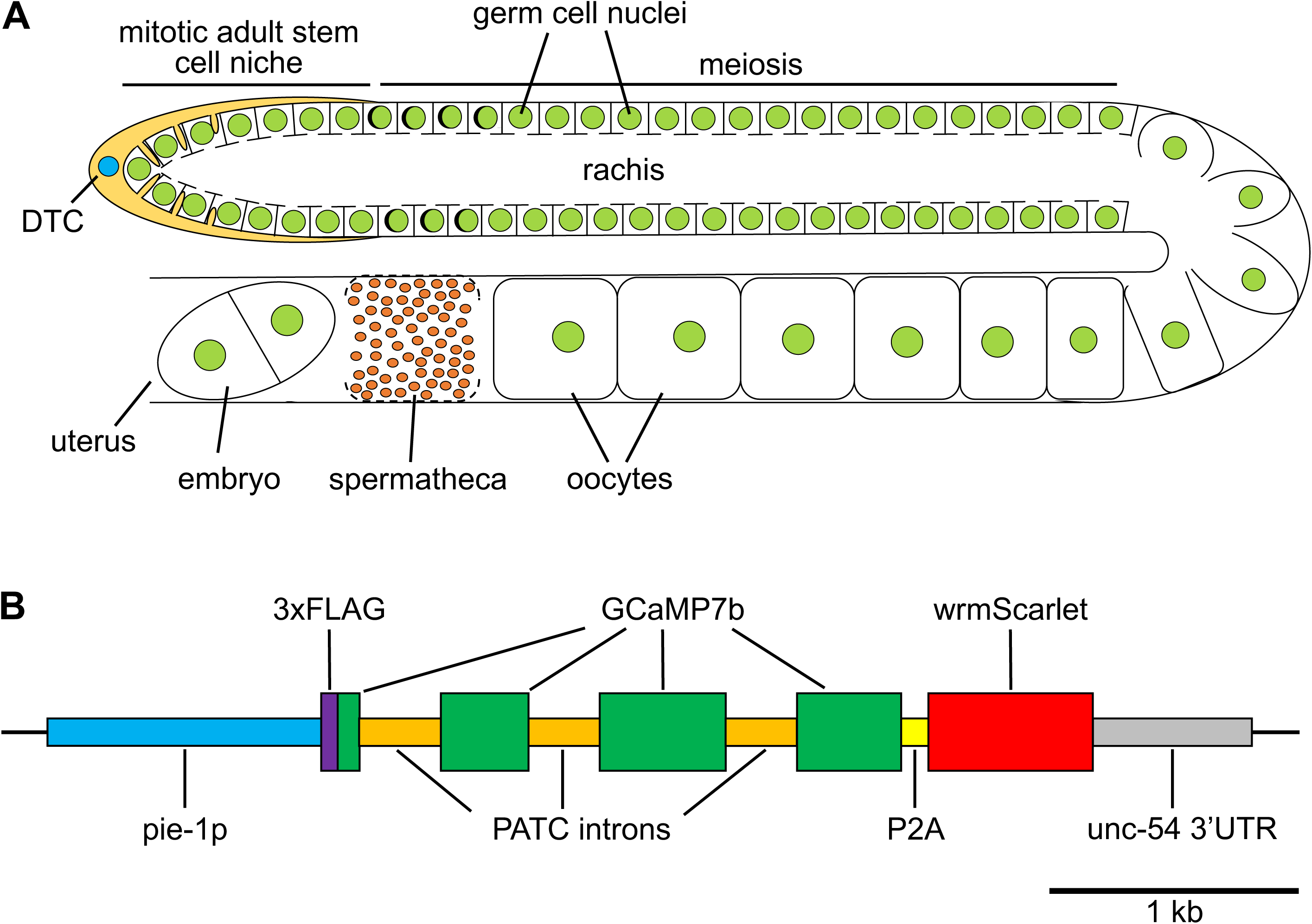
Diagrams of the *C. elegans* adult hermaphrodite gonad and GCaMP7b::wrmScarlet genomic insertion. **A**) Diagram of one of the two U-shaped gonads in an adult hermaphrodite. The distal tip cell (DTC), which is a somatic gonadal cell that creates the mitotic adult stem cell niche, is labeled. Germ cell nuclei are in green. Note the opening of germ cells to the common cytoplasm, termed the rachis. Sperm are shown in red within the spermatheca somatic gonadal structure. B) Diagram of the *pie-1p*::3xFLAG::GCaMP7b::P2A::wrmScarlet transgenic insertion in the SKI LODGE locus on chromosome III.

In adult hermaphrodites, mitotic germ stem cell proliferation is confined to the most distal region of the germ line in an adult stem cell niche defined by the projections of the distal tip cell (DTC) (Fig. 1A).^2^ The DTC signals via GLP-1/Notch to maintain germ cells in a mitotic state. As cells leave the DTC-defined stem cell niche, reduced Notch signaling and the action of pro-meiotic RNA regulatory pathways promote meiotic entry and differentiation.^1,3^ Once germ cells in adult hermaphrodites enter meiosis, they begin a regulatory program to become oocytes.

The *C. elegans* germ line is a syncytium in which germ cells are connected to a common cytoplasm through openings in their plasma membrane (Fig. 1A).^4^ This syncytium allows the germ cells to produce maternal product (mRNA and protein) that is transferred to the developing oocytes to allow rapid cell divisions in the early embryo. There does not appear to be mixing of individual germ cell cytoplasm because mitosis is not synchronous, whereas it would be synchronous if activated mitotic cyclin/CDK complexes transferred between mitotic and non-mitotic cells.^5^

In the wild, *C. elegans* generally spend most of their lives foraging for bacteria, their food source, in a nutrient-deprived state.^6^ When *C. elegans* hermaphrodites encounter rotting vegetation full of bacteria, they respond by increasing their rate of germ cell proliferation to rapidly produce progeny. Nematodes that can generate progeny faster have a clear evolutionary advantage. This is particularly the case because an adult nematode can produce ∼300 progeny in 3 days, and each of those progenies can produce ∼300 progeny in ∼6 days, quickly depleting the limited food supply.

The rate of generating germ cells in the mitotic zone is modulated by signaling pathways in response to environmental conditions (mainly the presence or absence of bacteria). Germ cell proliferation is promoted in response to the availability of food through insulin signaling via the insulin receptor binding of specific insulin ligands (INS-3 and INS-33),^7^ TGF-β signaling through the TGF-β receptor binding TGF-β,^8^ as well as cell intrinsic assessment of nutrient availability that involves S6 kinase and the TOR/RAPTOR pathway.^9^ Loss of these signaling pathways cause a decrease in germ cell proliferation, and starvation causes a cessation of proliferation.

In the *C. elegans* hermaphrodite gonad, calcium (Ca^2+^) plays several roles associated with fertilization and the initiation of embryogenesis. Major sperm protein, secreted by sperm, binds receptors on the proximal oocyte, including the Eph receptor VAB-1, and promotes meiotic maturation via Ca^2+^-dependent pathways, including calmodulin-dependent protein kinase II.^10^ However, an increase in the level of Ca^2+^ in the proximal oocyte has not been reported.^11^

Fertilization of an oocyte in the spermatheca initiates a wave of Ca^2+^ at the site of sperm entry that propagates across the oocyte.^12^ This fertilization Ca^2+^ wave prevents polyspermy and initiates embryogenesis, and is present during egg activation for all animal species examined.^13^ Oocyte entry into the spermatheca initiates a series of IP3-dependent Ca^2+^ oscillations in the spermatheca that lead to constrictions that move the zygote into the uterus.^14^

Here we report the use of a ratiometric Ca^2+^ reporter to allow measurements of basal (steady-state) levels of cytoplasmic Ca^2+^ in the germline. To date, changes in Ca^2+^ levels have not been linked to proliferation in *C. elegans*. We show that in the germ line, the basal level of cytoplasmic Ca^2+^ correlates with the proliferation rate of mitotic germ cells. Moreover, altering basal Ca^2+^ levels alters the proliferation rate of germ cells, indicating that basal Ca^2+^ levels can regulate proliferation.

## Results

### A ratiometric germline cytoplasmic Ca^2+^ reporter

In order to assess the relative levels of Ca^2+^ in the *C. elegans* germ line, we created an integrated genetically-encoded ratiometric Ca^2+^ reporter that is expressed in germ cells. We utilized the SKI LODGE system in which tissue-specific reporters are placed in the genome upstream of a CRISPR targeting sequence from the *dpy-10* gene.^15^ We used two rounds of CRISPR/Cas9 genome editing to place the Ca^2+^ reporter GCaMP7b^16^ and wrmScarlet (separated by a P2A ribosome skipping sequence) into the SKI LODGE site on chromosome III that contains the germline-specific promoter *pie-1p*. The GCaMP7b sequence was modified by optimizing the codons for *C. elegans* and introducing three PATC-enriched introns, which promote expression of transgenes in the germ line (Fig. 1).^17^ The transgene is referred to here as pGCS for *<u>p</u>ie-1p*::<u>GC</u>aMP7::P2A::wrm<u>S</u>carlet. During translation of pGCS mRNA, the newly synthesized GCaMP7b protein is released from the ribosome as it encounters the P2A ribosome skipping sequence, and then the ribosome translates a separate wrmScarlet protein. The wrmScarlet functions as an expression control, so that the ratio of GCaMp7b signal to wrmScarlet signal provides a ratiometric assessment of Ca^2+^ levels. To ensure that conditions of the microscope (e.g., confocal laser strength) did not affect quantitative comparisons, we always ran control samples at the same session and standardized the experimental results relative to the control samples.

### Ca^2+^ levels are altered during the sexual development of the hermaphrodite germ line

The GCaMP7b expressed from the pGCS transgene appears to be largely excluded from nuclei, and is therefore a cytoplasm-specific marker of Ca^2+^ levels (Fig. 2). The GCaMP7b/wrmScarlet ratio is spatially uniform throughout the gonad from the L1 through the L3 larval stages. The L4 larval stage can be subdivided into nine stages (L4.1 to L4.9) based on the morphology of the vulva.^18^ Beginning at approximately the L4.5 stage, several contiguous germ cells with high Ca^2+^ levels are observed near the bend in the gonad (Fig. 2B). The elevated Ca^2+^ levels in these cells continue until the L4.8 or L4.9 stage (Fig. 2C,D), at which time a lower level of increased Ca^2+^ is observed in more cells in the proximal region of the germ line that continues into the young-adult stage (Fig. 2E,F). The switch from only a few germ cells with very elevated Ca^2+^ levels to more cells with less elevated Ca^2+^ levels occurs in the region that gives rise to oocytes, which have also have elevated Ca^2+^ levels (Fig. 2D–H).

**Figure 2.**
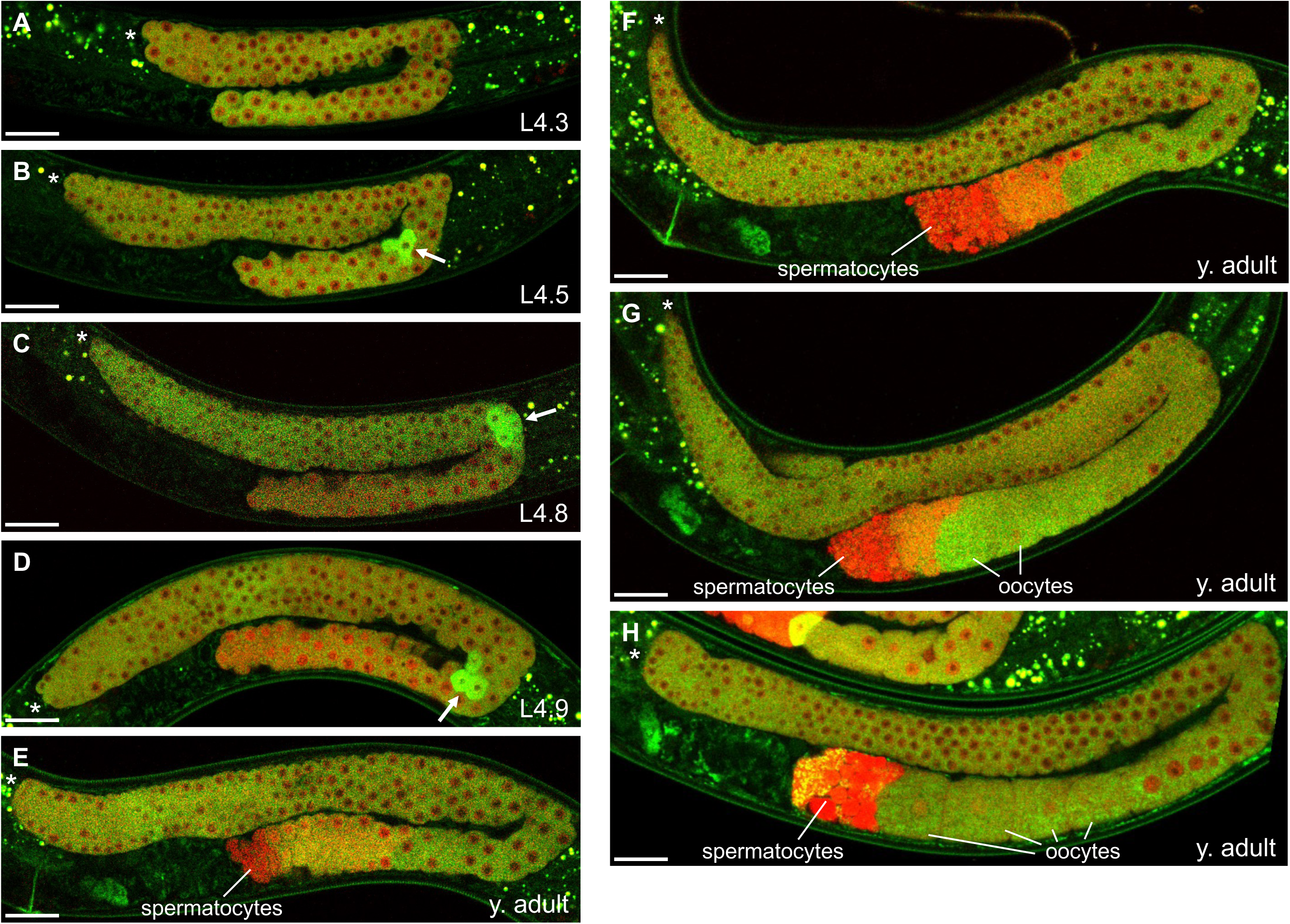
Ca^2+^ levels during hermaphrodite sexual development in L4 and young adult stages. GCaMP7b (green) and wrmScarlet (red) overlay of confocal images of hermaphrodites of the indicated ages, presented from younger (**A**) to older (**H**) animals. L4-stage larvae are denoted by their vulval morphology stage. Young adults (without eggs) are ordered based on the development of their oocyte region. Asterisks indicate the distal end of the gonad. Arrows indicate germ cells with high Ca^2+^ in the bend region of late L4 larvae. Oocytes and spermatocyte region are marked. Slight sections of (**E**) and (**H**) are cut off at the edge because the image was at an angle before cropping. Scale bars, 20 µm. Grayscale images for the two separate channels are shown in Supplemental Material Fig. S1.

In contrast to the region producing oocytes, the proximal region that gives rise to sperm has a reduced GCaMP7b/wrmScarlet ratio beginning around the L4.9 stage and then continues to have progressively lower GCaMP7b/wrmScarlet ratios in young adults. The spermatocyte GCaMP7b/wrmScarlet region is pushed more proximally by the developing oocytes, which have higher GCaMP7b/wrmScarlet ratios (Fig. 2F–G). There is an accumulation of wrmScarlet in the spermatocyte region, but also a loss of GCaMP7b signal (Suppl. Fig. S1). The most Ca^2+^-depleted objects in the spermatocyte region correspond to the residual bodies that remain after sperm formation (Fig. 2H; Suppl. Fig. S2). Sperm also have reduced Ca^2+^ levels, but the levels are higher than for the residual bodies (Suppl. Fig. S2).

### Reductions in germ cell proliferation reduce basal Ca^2+^ levels

We investigated whether conditions that reduce germ cell proliferation are correlated with changes in basal cytoplasmic Ca^2+^ levels in the germ line, particularly in the distal mitotic zone. We analyzed four conditions that are known to decrease mitotic germ cell proliferation.

Insulin signaling is known to promote germ cell proliferation.^7^ A null mutation of the insulin ligand *ins-3*(*ok2488*) significantly reduces germ cell proliferation (the mitotic index is ∼40% lower than in wild type).^7^ We found that the level of Ca^2+^ in *ins-3*(*ok2488*); pGCS L4-stage mutants was reduced 24.5% relative to wild-type levels (Fig. 3A).

**Figure 3.**
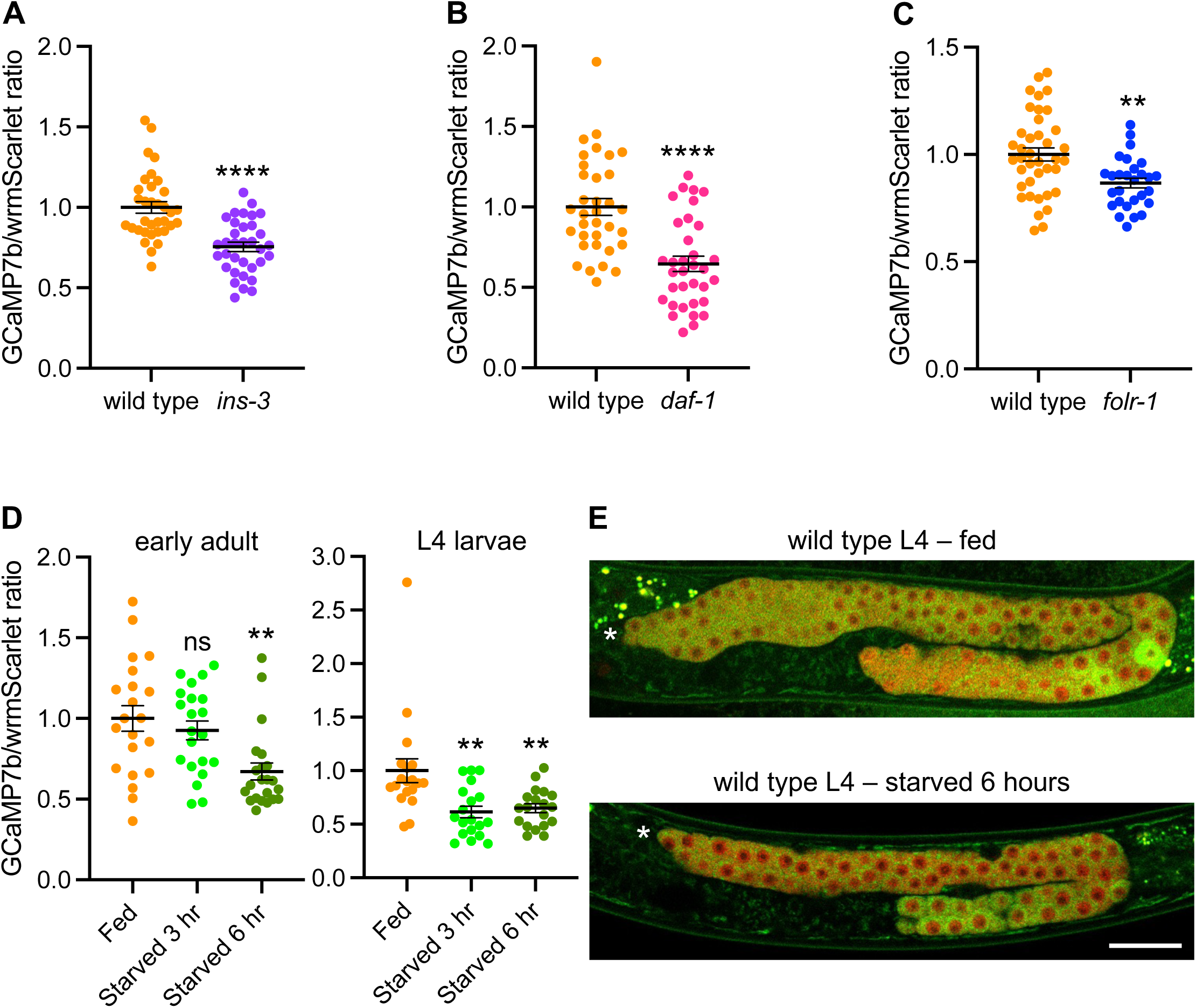
Ca^2+^ levels in animals with reduced germ cell proliferation. Graphs of the GCaMP7b/wrmScarlet ratios in L4-stage *ins-3*(*ok2488*); pGCS (**A**), early adult *daf-1*(*m40*ts); pGCS at 25°C for 1 day (**B**), and L4-stage *folr-1*(*ek44*); pGCS (**C**) relative to comparably staged wild-type pGCS animals. **D**) The GCaMP7b/wrmScarlet ratios in wild-type pGCS early adults (with 1 row of eggs of approximately 6-10 eggs) or L4-stage larvae starved for 3 to 6 hr or fed. All GCaMP7b/wrmScarlet ratio graphs are standardized to the control (wild type or fed) levels set to 1.0. **E**) Confocal images of wild-type pGCS L4-stage larvae that were fed or starved for 6 hr, with GCaMP7b (green) and wrmScarlet (red) overlay. Note the punctate GCaMP7b signal in the distal side of the germ line for the 6 hr starved L4 larva. Asterisks indicate the distal end of the gonad. Scale bar, 20 µm. Statistical analysis for (**A**)–(**C**) was with unpaired two-tailed Student’s t-test. Statistical analysis for (**D**) was with one-way ANOVA with Dunnett multiple comparisons. For all figures, error bars show the mean and standard error of the mean (SEM). For all figures, *p-value < 0.05; **p-value<0.01; ***p-value<0.001; ****p-value<0.0001. Grayscale images for each channel in (**E**) are shown in Supplemental Fig. S3.

TGF-β signaling is also known to promote germ cell proliferation.^8^ *daf-1*(*m40*ts) is a temperature sensitive mutation of the TGF-β type I receptor. Switching *daf-1*(*m40*ts) L4 larvae from 16°C (the permissive temperature) to 25°C (the restrictive temperature) for one day reduces the number of cells in the germline mitotic zone of the resulting early adults by 38.5%.^8^ We repeated this with *daf-1*(*m40*ts); pGCS mutants and found that the level of Ca^2+^ was reduced 35.5% relative to wild-type levels (Fig. 3B).

Loss of FOLR1-mediated folate signaling produces germ cells that are unable to increase their proliferation rate in response to folate obtained from ingested bacteria.^19^ We found that *folr-1*(*ek44*) null mutant L4 larvae have a 13% reduction in basal Ca^2+^ levels (Fig. 3C). Thus, inactivation of three different signaling pathways that result in reduced germ cell proliferation rates were each correlated with reduced levels of basal cytoplasmic Ca^2+^.

Starvation significantly reduces germ cell proliferation in *C. elegans*.^20-22^ We tested the effect of starvation on germline Ca^2+^ levels. Early adults (approximately one day old with one row of 10 eggs or less) were starved for 3 or 6 hr. At 3 hr, there was not a significant decrease in the germline basal Ca^2+^ levels (Fig. 3D). However, by 6 hr, the basal Ca^2+^ levels were reduced 33% relative to fed animals (Fig. 3D). When L4-stage animals were starved, they showed a more rapid reduction in basal cytoplasmic Ca^2+^ levels. The basal level of Ca^2+^ in 3 hr starved L4 larvae was reduced 38.5% relative to fed animals, and this reduced level was also observed at 6 hr of starvation (a reduction of 35% relative to fed animals) (Fig. 3D).

The cytoplasmic pattern of GCaMP7b germ cell signal in starved animals was different than in fed wild type or the mutants with reduced germ cell proliferation. In addition to the reduction in GCaMP7b signal relative to wrmScarlet, the remaining GCaMP7b signal was observed in puncta that often localized to the cell periphery (rather than the more uniform signal observed throughout the cytoplasm) (Fig. 3E). Overall, four different methods of reducing germ cell proliferation all led to decreases in basal cytoplasmic Ca^2+^ levels.

### Tumorous overproliferation is associated with increased basal Ca^2+^ levels

To assess whether increased proliferation rates lead to an increase in Ca^2+^ levels in hermaphrodites, we analyzed proximal tumors in the *cki-2*(*ok2105*) and *daf-16*(*mu86*); pGCS strain. CKI-2 is a CDK inhibitor and DAF-16 is a FOXO transcription factor, both of which act to inhibit germ cell proliferation.^7,23^ We found that shifting the *cki-2*(*ok2105*) and *daf-16*(*mu86*); pGCS strain from 16°C to 25°C produces proximal germline tumors (Fig. 4A). Animals were shifted for two days to 25°C, and animals with visible tumors (the majority of adults) were analyzed for the level of Ca^2+^. Significantly, the proximal tumors in *cki-2*; *daf-16*(*mu86*); pGCS animals had elevated levels of Ca^2+^ relative to the mid region or distal region of the gonads (Fig. 4B). The GCaMP7b/wrmScarlet ratio in *cki-2*; *daf-16*(*mu86*); pGCS proximal tumors were 2.7- to 2.9-fold higher than in middle (distal to the bend) and distal gonad regions of the same animals (Fig. 4B). The GCaMP7b/wrmScarlet ratio of the proximal tumors were also almost 3-fold higher than in the proximal regions of wild-type pGCS animals (Fig. 4B). These results indicate that the over-proliferating proximal tumor cells have higher levels of basal cytoplasmic Ca^2+^ than non-tumorous germ cells.

**Figure 4.**
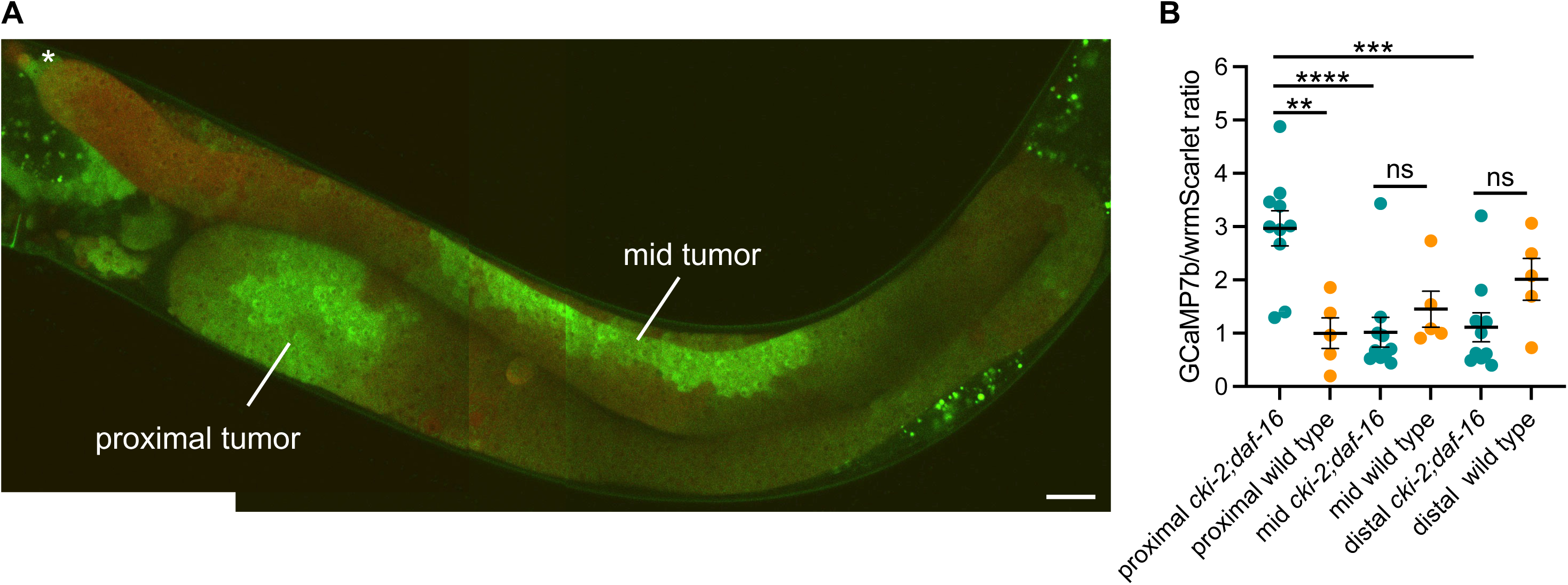
*cki-2*; *daf-16* mutants have increased Ca^2+^ in their proximal tumors. **A**) Composite image of GCaMP7b and wrmScarlet overlay of proximal tumors in a *cki-2*; *daf-16*; pGCS adult hermaphrodite grown at 25°C for 48 hr. Tumors are labeled. The distal tip of the gonad is marked with an asterisk. The focal plane varies slightly between the three individual images of the compound image. Scale bar, 20 µm. **B**) Graph of GCaMP7b/wrmScarlet ratios in the proximal region or proximal tumor, mid region, or distal mitotic zone of the germ line for *cki-2*; *daf-16*; pGCS and wild type; pGCS. Statistical analysis was with one-way ANOVA with Dunnett multiple comparisons. Grayscale images for each channel in (**A**) are shown as Supplemental Fig. S4.

### Reductions in basal Ca^2+^ levels reduce proliferation

So far, we have shown that conditions that decrease proliferation have lower basal levels of Ca^2+^, and conditions with increased proliferation have higher basal levels of Ca^2+^. We wanted to determine if Ca^2+^ levels were instructive for proliferation by reducing the level of basal germ cell Ca^2+^ and determining the effect on proliferation. To reduce cytoplasmic Ca^2+^ levels in the germ line, we inactivated the TRPM7-related Ca^2+^ channel GON-2.

The *gon-2* gene encodes a TRPM-related Ca^2+^ channel, whose closest human homolog is TRPM7.^24^ *gon-2* mutants were first identified based on temperature-sensitive mutants in which the somatic gonad precursors do not divide at the restrictive temperature to form the normal hermaphrodite gonad structure.^25^ A GON-2::GFP transgene demonstrates the expression of GON-2::GFP in germ cells, with an enrichment on the outer surface of the germ cells (Fig. 5A). Unlike the temperature sensitive mutations, *gon-2*(*ok465*) null homozygotes form the somatic gonad structure. We observed that *gon-2*(*ok465*); pGCS animals have a 42% reduction in the GCaMP7b/wrmScarlet ratio in the distal region of the gonad (Fig. 5B,C). Notably, *gon-2*(*ok465*) mutants have a severe reduction in the number of germ cells relative to wild type and are sterile (Fig. 5B).^26^ Late-stage L4 *gon-2*(*ok465*); pGCS homozygotes had only 30.9 ± 2.8 (SEM) germ cells (n = 14) on average in the entire region of the germ line that lies along the dorsal side. In contrast, there are 127.0 ± 5.9 germ cells (n = 13) in the mitotic zone region of L4-stage wild type; pGCS animals, with the mitotic zone encompassing just a portion of the dorsal side of the germ line. The reduction in *gon-2*(*ok465*) germ cell numbers relative to wild type is significant to a p-value < 0.0001.

**Figure 5.**
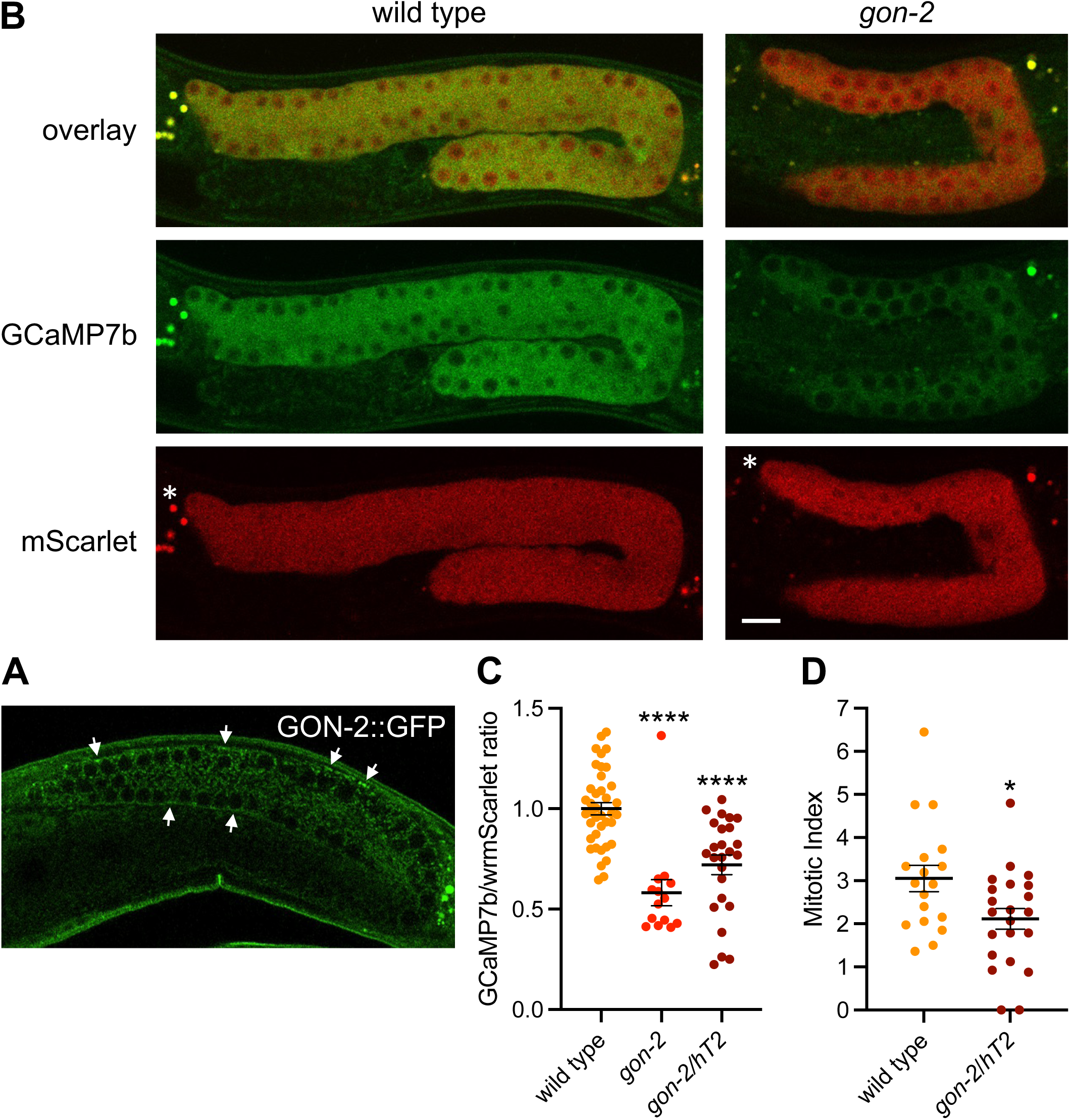
*gon-2* null mutants have reduced basal cytoplasmic Ca^2+^ levels and germ cell numbers. **A**) Confocal image showing expression of *gon-2p*::GON-2::GFP in the germ line of an adult hermaphrodite, with enrichment at the outer edge of the germ cells (arrows). Asterisks indicate the distal end of the gonad. **B**) Confocal images of L4-stage wild type and *gon-2*(*ok465*) animals with GCaMP7b and wrmScarlet presented separately and overlaid. **C**) Graph of the GCaMP7b/wrmScarlet ratio in *gon-2*(*ok465*) homozygotes and *gon-2*(*ok465*)/*hT2* heterozygotes. **D**) Graph of the number of mitotic cells per gonad in *gon-2*(*ok465*)/*hT2* heterozygotes and wild type. Statistical analysis for (**C**) was with one-way ANOVA with Dunnett multiple comparisons. Statistical analysis for (**D**) was with unpaired two-tailed Student’s t-test. Scale bar, 20 µm.

Potentially, the decrease in Ca^2+^ levels in *gon-2*(*ok465*) mutants arose from having too few germ cells. To assess this, we determined the level of Ca^2+^ in *gon-2*/*hT2* heterozygotes, which have relatively normal numbers of germ cells (124.2 ± 8.1 germ cells per mitotic zone in L4 larvae; n = 16) and are fertile. *gon-2*/*hT2* heterozygotes have a 28% lower level of Ca^2+^ in the distal mitotic zone relative to wild type (Fig. 5C). The reduction in Ca^2+^ in *gon-2*/*hT2* heterozygote animals correlates with a 31% lower mitotic index than in wild type (Fig. 5D). Therefore, *gon-2* heterozygotes with an overtly normal germ line also exhibit reduced germ cell basal Ca^2+^ levels, implying that a partial loss of GON-2 levels produces a partial reduction in cytoplasmic Ca^2+^ levels. Thus, the GON-2 Ca^2+^ channel is required to maintain basal cytoplasmic Ca^2+^ levels in germ cells, and the reduction in Ca^2+^ levels when GON-2 is inactivated is associated with decreased germ cell proliferation.

### Increases in basal Ca^2+^ levels increase proliferation

Having shown that decreases in basal cytoplasmic Ca^2+^ lead to reduced proliferation, we wanted to determine whether increases in Ca^2+^ levels would increase proliferation. To accomplish this, we focused on two essential regulators that remove cytoplasmic Ca^2+^: the SERCA channel, which transports cytoplasmic Ca^2+^ into the endoplasmic reticulum; and the Plasma Membrane Ca^2+^ ATPase (PMCA), which transports cytoplasmic Ca^2+^ out of the cell. There is only one SERCA gene in *C. elegans*, *sca-1*.^27^ There are three PMCA genes in *C. elegans*, *mca-1*, *mca-2*, and *mca-3*.^28^ Of these three genes, only *mca-3* is expressed to high level in the germ line.^29^ Partial inhibition of *sca-1* or *mca-3* are known to increase cytoplasmic Ca^2+^ levels in other *C. elegans* tissues.^27,30^

We observed that 10% *sca-1* RNAi (10% *sca-1* RNAi bacteria mixed with 90% control RNAi bacteria) increased basal cytoplasmic Ca^2+^ levels in distal germ cells by 20% relative to control RNAi in L4-stage larvae (Fig. 6A). 50% *mca-3* RNAi increased basal cytoplasmic Ca^2+^ levels in distal germ cells by 79% relative to control RNAi in L4 larvae (Fig. 6B).

**Figure 6.**
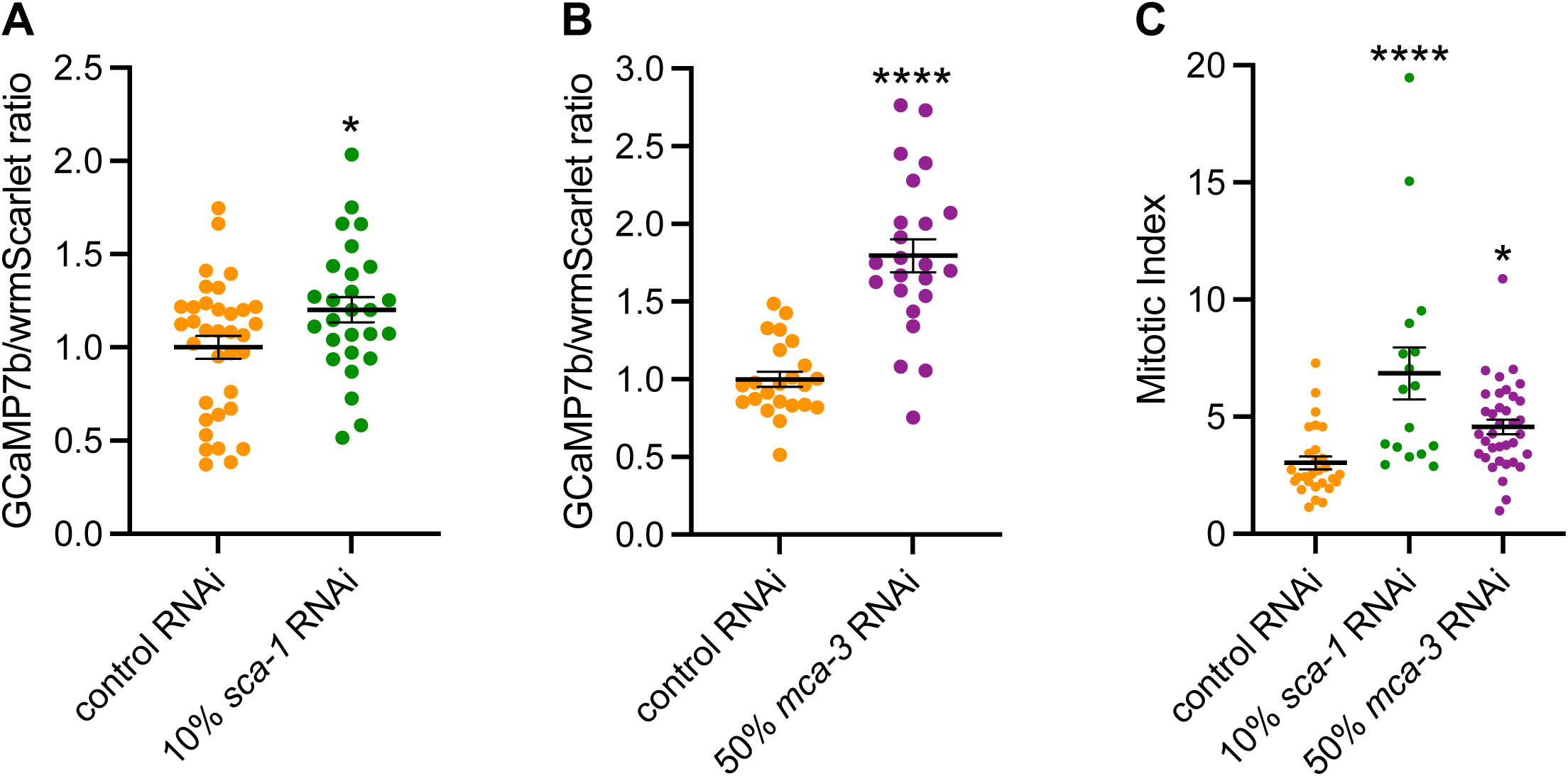
Increasing basal cytoplasmic Ca^2+^ increases germ cell proliferation. **A**) Graph of the GCaMP7b/wrmScarlet ratios in pGCS L4 larvae subject to 10% *sca-1* or control RNAi. **B**) Graph of GCaMP7b/wrmScarlet ratios in pGCS L4 larvae subject to 50% *mca-3* or control RNAi. **C**) Mitotic index of pGCS L4 larvae treated with control RNAi, 10% *sca-1* RNAi, or 50% *mca-3* RNAi. Statistical analysis for (**A**) and (**B**) was with unpaired two-tailed Student’s t-test. Statistical analysis for (**C**) was with one-way ANOVA with Dunnett multiple comparisons.

We analyzed proliferation in L4-stage larvae. 10% *sca-1* RNAi increased the mitotic index of germ cells in the mitotic zone two-fold relative to wild type (Fig. 6C). 50% *mca-3* RNAi increased the mitotic index by 50.5% relative to wild type (Fig. 6C). Thus, we conclude that increasing basal cytoplasmic Ca^2+^ levels through two different cellular mechanisms increases germ cell proliferation.

## Discussion

Here we provide the first report of the level of cytoplasmic Ca^2+^ in the *C. elegans* germ line using a stably integrated ratiometric Ca^2+^ reporter. The use of an integrated ratiometric Ca^2+^ reporter allows more accurate comparisons of the level of Ca^2+^ during development. Our analysis shows that there are changes in Ca^2+^ levels during sexual development with a reduction in the Ca^2+^ levels of spermatocytes and an increase in oocyte precursors. Notably, we observe that several cells in the bend region of mid-L4 larvae have elevated Ca^2+^. The germ cells with elevated Ca^2+^ are localized more proximally in late L4, and then the pattern switches from a few cells with very elevated Ca^2+^ levels to more cells with less elevated Ca^2+^ levels in the oocyte-forming region. The striking elevation of Ca^2+^ in only certain germ cells in mid-to-late L4 gonads suggests that they are developmentally different than the surrounding germ cells. Studies on the oocyte-expressed yolk receptor RME-2 reported initial expression of RME-2::GFP in the bend region.^31^ This suggests that the initial cells with elevated Ca^2+^ levels may be oocyte precursors.

We have observed that conditions that reduce the proliferation rates of germ cells correlate with reductions in basal cytoplasmic Ca^2+^ levels. These conditions include loss of insulin, TGF-β, and FOLR1 signaling, and starvation. Conversely, germline tumors, which have increased proliferation show elevated basal cytoplasmic Ca^2+^ levels. These correlations suggest either that basal Ca^2+^ levels are responsive to the proliferation rate (with Ca^2+^ levels decreasing or increasing in response to decreased or increased proliferation rates, respectively) and/or that Ca^2+^ levels are regulated by signaling events that also alter proliferation rates. These are not mutually-exclusive scenarios, and both could be operating in different physiological contexts.

Our data on starvation suggests that reductions in Ca^2+^ levels do not directly follow from a reduction in the rate of proliferation, but may instead respond to other cues, such as metabolism. In early adult hermaphrodites, basal Ca²⁺ levels did not change at 3 hr of starvation but were significantly reduced by 6 hr. In contrast, L4 larvae showed a significant reduction in basal Ca²⁺ at 3 hr (similar to 6 hr, suggesting a plateau). These results do not correlate with mitotic proliferation. Published work shows that in starved early adults the number of mitotic cells per gonad drops by about half within the first hour and is near zero by 3 hr (at which time there is no effect on basal Ca^2+^ levels).^22^ In contrast, in starved L4 larvae, mitotic cell numbers are reduced by less than 50% at 3 hr (yet there is a significant drop in Ca^2+^ levels).^22^ Therefore, changes in basal Ca²⁺ levels do not correlate with changes in proliferation.

A plausible hypothesis is that the decrease in Ca^2+^ during starvation is in response to metabolism or signaling associated with metabolism. Notably, adults have significantly more fat storage than L4 larvae,^32^ with the triacylglyceride-to-protein ratio twice as high in early adults compared to L4 larvae.^33^ Thus, early adults have substantially more fat stores that are likely to delay the effects of starvation on metabolism relative to L4 larvae.

We observed that decreasing Ca^2+^ levels, by inactivating the GON-2 Ca^2+^ channel, decreases proliferation. Conversely, increasing Ca^2+^ levels, by partially inactivating the SERCA and PMCA mechanisms for removing cytoplasmic Ca^2+^, led to increases in germ cell proliferation. This suggests that the level of basal cytoplasmic Ca^2+^ is instructive for the rate of germ cell proliferation. This data supports a model in which the level of basal cytoplasmic Ca^2+^ acts as a rheostat to modulate the rate of germ cell proliferation.

Our study suggests that the TRPM7-channel GON-2, which is expressed in the germ line, is a central regulator of basal cytoplasmic Ca^2+^ levels. Inactivation of *gon-2* leads to a 42% reduction in the level of basal cytoplasmic Ca^2+^ in the distal region of the gonad. This significant reduction in basal cytoplasmic Ca^2+^ levels is associated with very few germ cells and sterility. That the basal level of Ca^2+^ in germ cells only goes down 42% suggests that other Ca^2+^ channels/transporters also function to transport Ca^2+^ into the germ line, but that these other channels/transporters cannot compensate for a complete lack to GON-2 to bring basal Ca^2+^ to normal physiological levels.

It is known that transient increases in Ca^2+^ that are associated with cell signaling can promote cell cycle progression in mammalian cells.^34^ Transient changes in Ca^2+^ linked to signaling are also associated with proliferation in the invertebrates *Drosophila melanogaster* and Planaria.^35,36^ While there are hundreds of studies on transient Ca^2+^ increases in response to signaling leading to increased cell proliferation, there are relatively few studies that demonstrate a correlation between proliferation and basal Ca^2+^ levels. Cultured primary human or rat pulmonary artery smooth muscle cells in culture have ∼2-fold higher basal cytoplasmic Ca^2+^ level when proliferating in the presence of serum than when quiescent in serum-deprived conditions.^37,38^ Additionally, raising the level of basal cytoplasmic Ca^2+^ by inhibiting store-operated Ca^2+^ entry in a mouse myoblast cell line increases proliferation.^39^ While there are a few examples with mammalian cells that link basal cytoplasmic Ca^2+^ levels to proliferation, our study shows that basal Ca^2+^ levels regulate proliferation in a tissue in an intact animal under physiological conditions. Here, we have found that normal physiological contexts of food intake, and cell signaling pathways regulate the basal level of cytoplasmic Ca^2+^ to affect the rate of germ cell proliferation. There are no other published studies linking Ca^2+^ levels (basal or transient) and cell proliferation in *C. elegans*.

Our study has thus identified that the basal cytoplasmic level of Ca^2+^ regulates cell proliferation in germ cells, and changes in basal Ca^2+^ levels are associated with multiple pathways that regulate germ cell proliferation. It will be interesting in future experiments to determine how different signaling pathways intersect with or regulate basal cytoplasmic Ca^2+^ levels to control the proliferation of *C. elegans* germ stem cells.

## Methods

### C. elegans strains and culture methods

*C. elegans* strains were cultured according to established methods.^40^ The following *C. elegans* strains were used. N2: wild-type Bristol strain, WBM1119: *wbmIs60*[*pie-1p*::3XFLAG::*dpy-10* crRNA::*unc-54* 3’UTR] (III:7007600), ET656: *ekIs32*[*pie-1p*::3XFLAG::GCaMP7b::*wrmScarlet*::*unc-54* 3’UTR], ET708: *ins-3*(*ok2488*) II; *ekIs32* III, ET709: *daf-1*(*m40*ts) IV; *ekIs3* III, ET670: *cki-2*(*ok2105*) II; *daf-16(mu86)* I; *ekIs32* III, ET666: *folr-1*(*ek44*) X; *ekIs32* III; ET678: *gon-2*(*q388*)ts I; *ekIs32*; and EJ938: *gon-2*(*q388*) I; *dxIs1*[*gon-2p*::GON-2::GFP], the kind gift of Eric Lambie.

The pGCS signal is not readily visible with a stereomicroscope equipped with fluorescence, but is easily observed with a confocal microscope, where z-sections can separate the green germline signal from the green autofluorescence of the intestine. Strains with the pGCS transgene spontaneously lose expression over time, potentially due to negative selective pressure from expression of GCaMP7b. To counteract this, actively growing reporter strains were maintained by weekly selection for animals with high levels of wrmScarlet expression. Strains that have lost high-level reporter expression can have their expression restored by sequentially selecting higher levels of expression in subsequent generations.

For selection of animals with higher pGCS expression, approximately 12 animals were put on microscope slides with agar pads in 6 µl of M9 buffer.^40^ The level of wrmScarlet was assessed with a compound fluorescence microscope. Animals with higher expression level were recovered by sliding the coverslip and picking up the worms with a curved platinum-iridium wire worm pick with bacteria.

Starvation was carried out by washing animals of the selected developmental stage six times in M9 buffer^40^ using polystyrene 15 ml tubes (Falcon). The animals were then placed on M9 agarose plates, containing M9 buffer with 15 g/ml of agarose (Sigma), and starved for the specified time.

To obtain proximal tumors in *cki-2(ok2105)*; *daf-16(mu86)*; pGCS strains; animals were shifted from 16°C to 25°C for 2 days. Animals with obvious tumors (encompassing the majority of adults) were selected for analysis.

### RNAi

RNAi was carried out with feeding-RNAi bacterial strains from the Ahringer library.^41^ RNAi bacteria were grown (from an overnight culture) in 2xYT media plus 100 µg/ml carbenicillin to an optical density at wavelength 600 nm (OD_600_) of between 0.4 and 0.6, at which time 1 mM IPTG (Gold Biotechnology) was added, and the culture was grown for 6 hr. The bacteria were concentrated by centrifugation and resuspension in a smaller volume of LB media with 100 µg/ml carbenicillin and 1 mM IPTG, and plated onto LB agar plates with 100 µg/ml carbenicillin and 1 mM IPTG. L4 stage hermaphrodites were placed on RNAi and their progeny were analyzed.

### Generation of the pie-1p::GCaMP7b::P2A::wrmScarlet integrated reporter

The GCaMP7b sequence with three PATC introns and 120 bp flanking sequences, for insertion into the *pie-1p*::*dpy-10* crRNA SKI LODGE locus in strain WBM1119, was synthesized by BioMatik Inc. Recombinant HIS6-Cas9 protein was biochemically isolated by our lab as described,^26^ using plasmid pHO4d-Cas9.^42^ The *dpy-10* crRNA, sequence GCUACCAUAGGCACCACGAG, was as described,^15^ and synthesized by Synthego. CRISPR/Cas9 was assembled in vitro according to Synthego instructions with 2.3 µM Cas9 and 2.4 µM annealed crRNA and tracrRNA (Synthego). The rescue DNA construct was a ssDNA–dsDNA–ssDNA annealing of the GCaMP7b sequence with the full GCaMP7b plus 120 bp flanking sequences so that the 120 ntd flanking sequences were ssDNA and the GCaMP7b sequence was dsDNA. The annealing procedure for the ss–ds–ss DNA rescue construct was as described.^43^ The injection mix included 2.3 µM Cas9 and 2.4 µM annealed crRNA and tracrRNA and 118 ng/µl rescue DNA. Gravid one-day old wild-type adults were injected in their gonads with the CRISPR/Cas9 rescue mix. Dpy progeny (due to CRISPR/Cas9 inactivation of the endogenous *dpy-10* locus) were isolated and screened for the insertion using PCR. The *dpy-10* mutation was eliminated by outcrossing the strain.

The insertion of the P2A::wrmScarlet sequence into the *pie-1p*::GCaMP7b SKI LODGE site was undertaken using the same procedure. The CRISPR crRNA sequence at the 3’ end of the inserted GCaMP7b coding sequence was GGGCGCGAGATGTTACTTGG, and was synthesized by Synthego. The P2A::wrmScarlet DNA sequence was synthesized by Azenta. A ss–ds–ss DNA rescue construct for P2A::wrmScarlet was generated and included in the injection mix as described above. Dpy progeny were isolated and screened for insertions by PCR. The strain with the P2A::wrmScarlet insertion was outcrossed several times.

### Microscopy

Non-confocal fluorescence and DIC images were obtained with a Zeiss Axioskop microscope, with images captured with either a Tucsen Dhyana 400 BSI V2 sCMOS camera using Micromanager software (version 1.4.23) or with a Hamamatsu ORCA-ER CCD camera using OpenLab 5.02. software (Improvision). Confocal images for the analysis of GCaMP7b–wrmScarlet were taken on a Zeiss LSM 880 confocal microscope, captured with Zen Blue software (Zeiss). The image for GON-2::GFP expression in the germline was obtained using an Andor Dragonfly Spinning Disk Confocal Microscope. Images were processed with Adobe Photoshop (version 22.0.0) and Fiji software (ImageJ version 2.1.0/1.53s). Matched fluorescence images to be used for level comparisons were processed identically and did not include gamma adjustments. For GCaMP7b/wrmScarlet ratios, control wild-type animals were always imaged at the same session, and the control GCaMP7b/wrmScarlet ratios were used to standardize the experimental ratios.

For figure images that are to be quantitatively compared, the level of wrmScarlet signal was used to standardize the intensity of GCaMP7b and wrmScarlet signals between the images to account for differences in expression level between animals. To accomplish this, the mean level of wrmScarlet signal for the entire distal-side of the germ line was determined. The ratio was determined for the mean wrmScarlet level in the higher-expression image divided by the mean wrmScarlet level in the lower-expression image. That ratio was used to increase the signal of both wrmScarlet and GCaMP7b channels in the images with lower wrmScarlet levels using the Fiji/ImageJ multiply function. Once the level of signal for the lower-expression image was raised to the level for the higher-expression image, the images were leveled to the same extent.

### Statistics

Every experimental result was repeated with comparable results with at least two biological replicates that were carried out on different days. Statistical analysis between two samples used unpaired two-tailed Student’s t-test (Figs 3A–C, 5D, 6A,B). Statistical analysis between multiple samples used one-way ANOVA with Dunnett for multiple comparisons (Figs 3D, 4B, 5C, 6D). Results are reported as mean ± SEM (standard error of the mean). For all figures, *p-value < 0.05; **p-value<0.01; ***p-value<0.001; ****p-value<0.0001.

## Acknowledgements

Some strains were provided by the CGC, which is funded by NIH Office of Research Infrastructure Programs (P40 OD010440). We acknowledge the assistance of the Biomedical Microscopy Core at the University of Georgia with imaging. This research was funded by a grant from the NIH National Institute of General Medical Sciences (NIGMS), R01GM134359 (ETK). We thank Dr. Eric Lambie for reagents.

## Supplementary Figure Legends

**Supplementary Figure S1.**
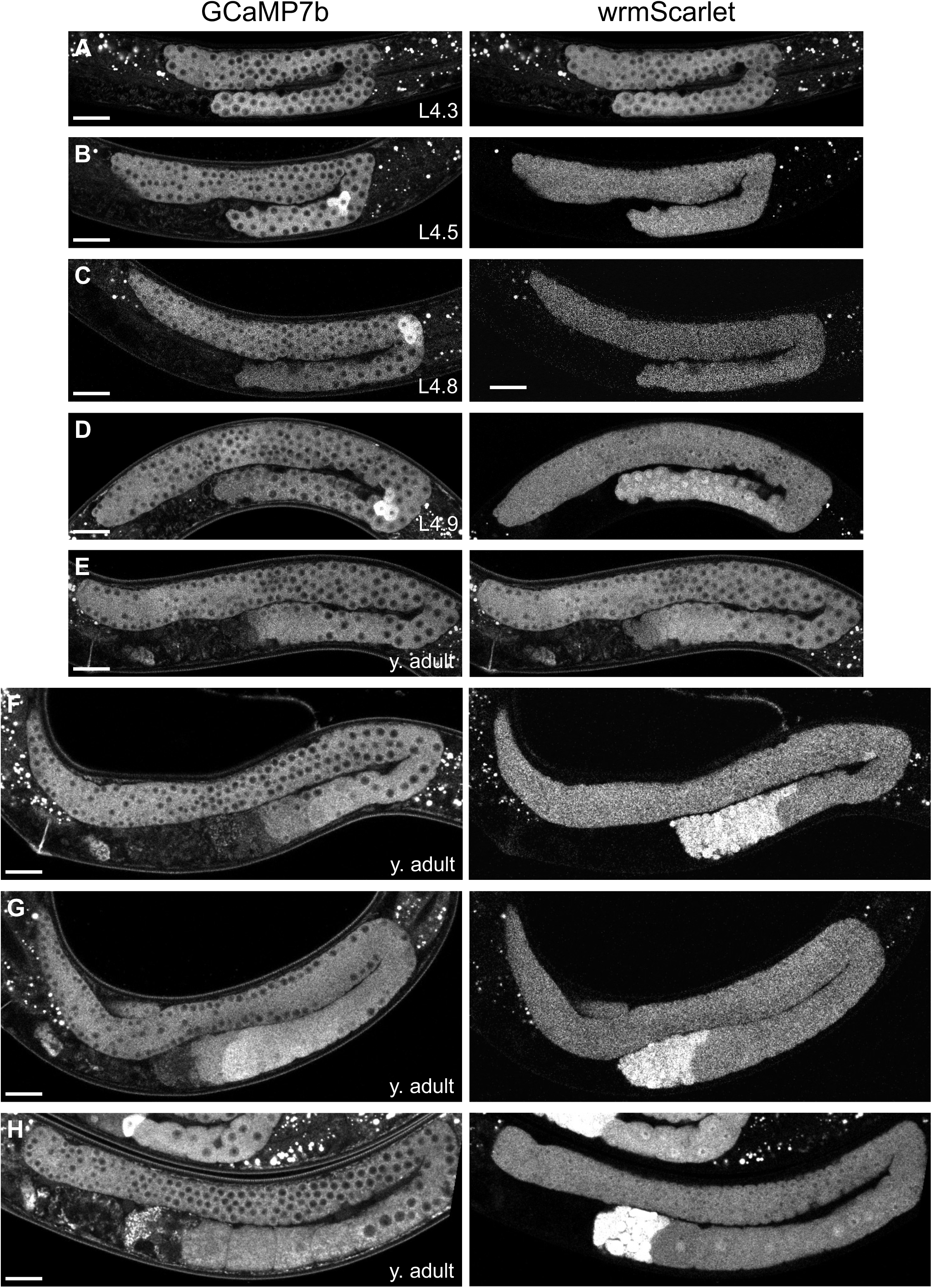
Single channel grayscale images of GCaMP7b and wrmScarlet for the overlaid images in Figure 2. Scale bars, 20 µm.

**Supplementary Figure S2.**
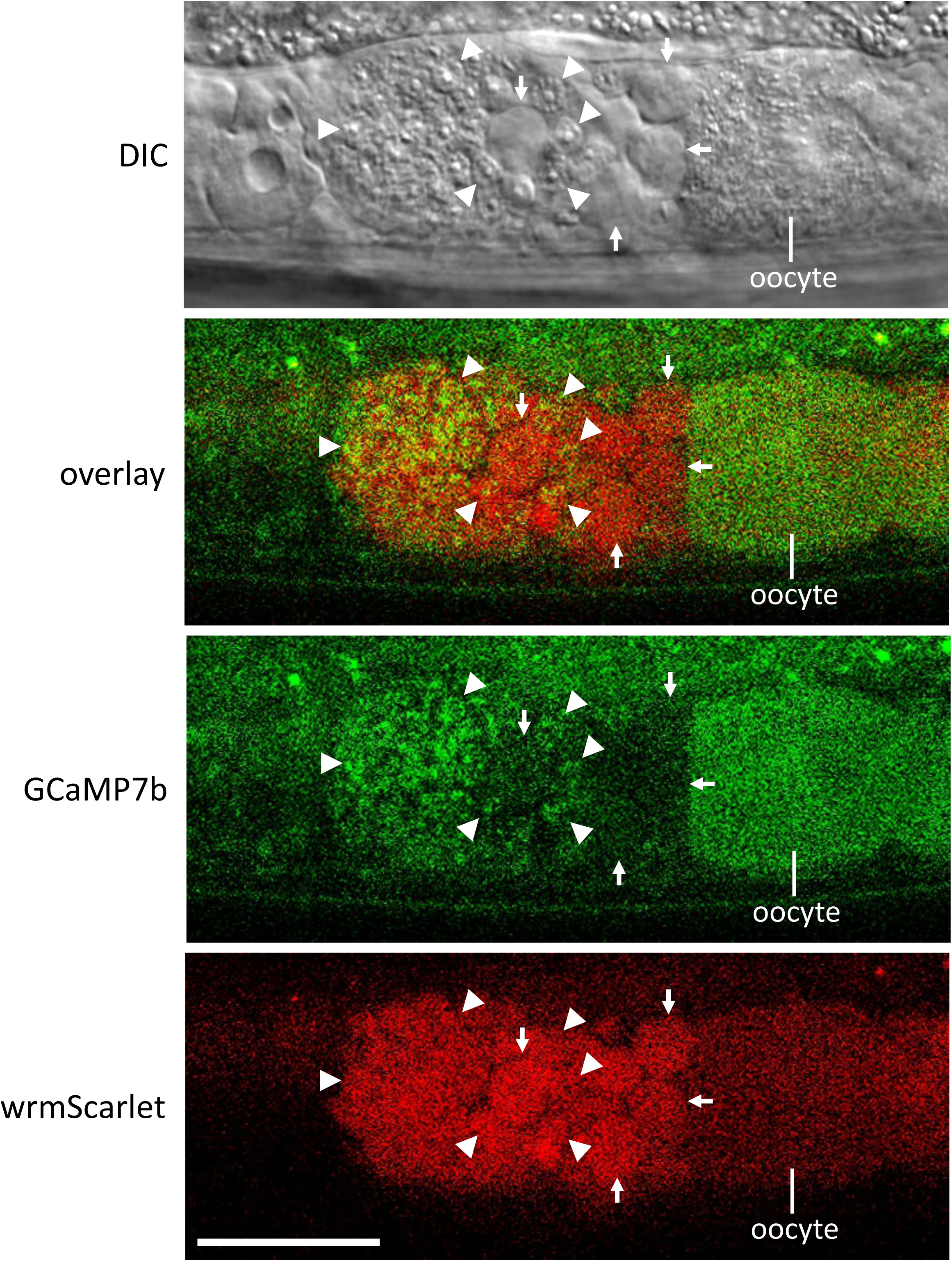
Sperm and proximal oocyte of wild type; pGCS young adult hermaphrodite. Confocal image of DIC, GCaMP7b, wrmScarlet, and overlay. Arrowheads, sperm. Arrows, residual bodies. Scale bar, 20 µm.

**Supplementary Figure S3.**
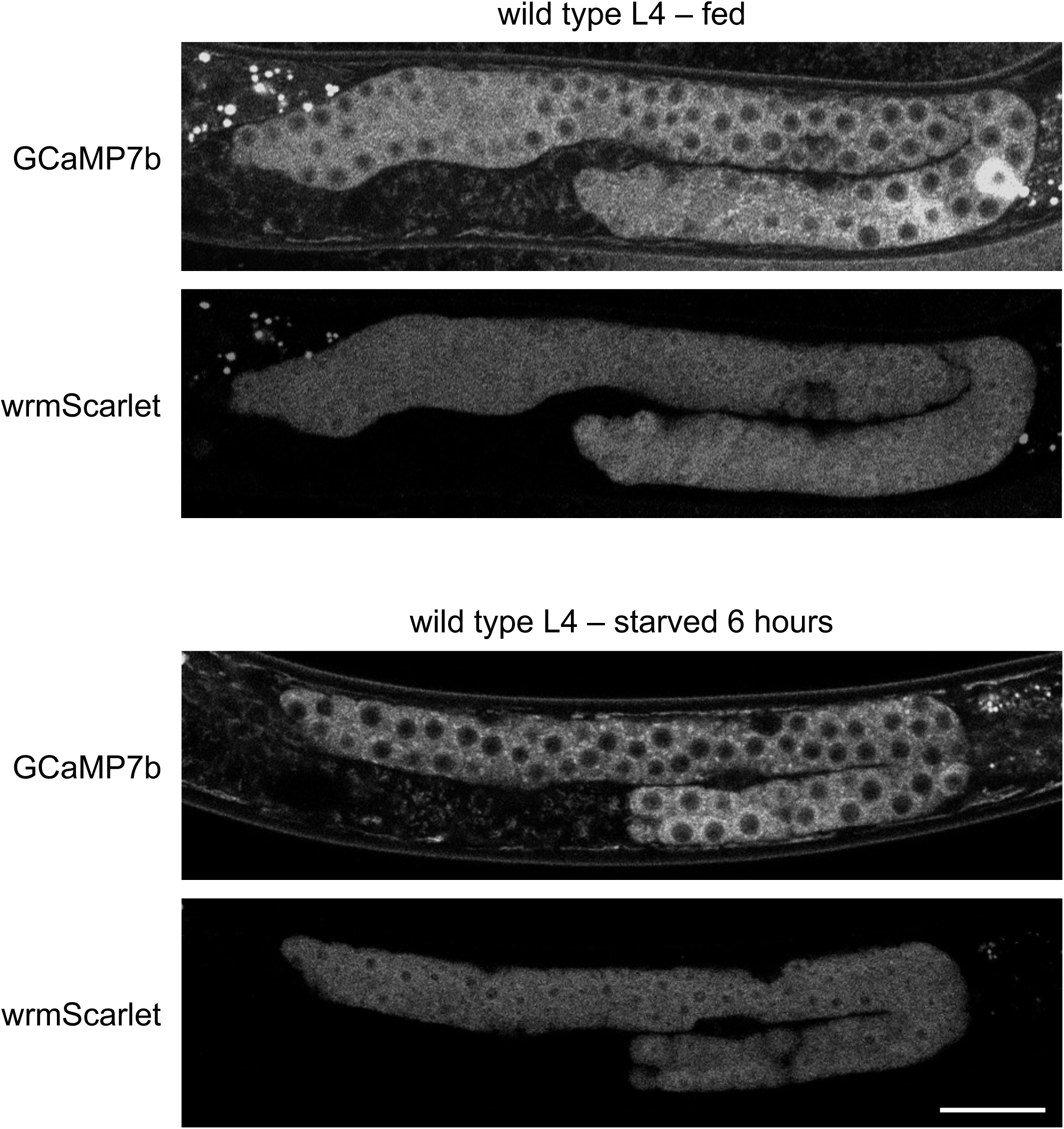
Single channel grayscale images of GCaMP7b and wrmScarlet for the overlaid images in Figure 3E. Scale bar, 20 µm.

**Supplementary Figure S4.**
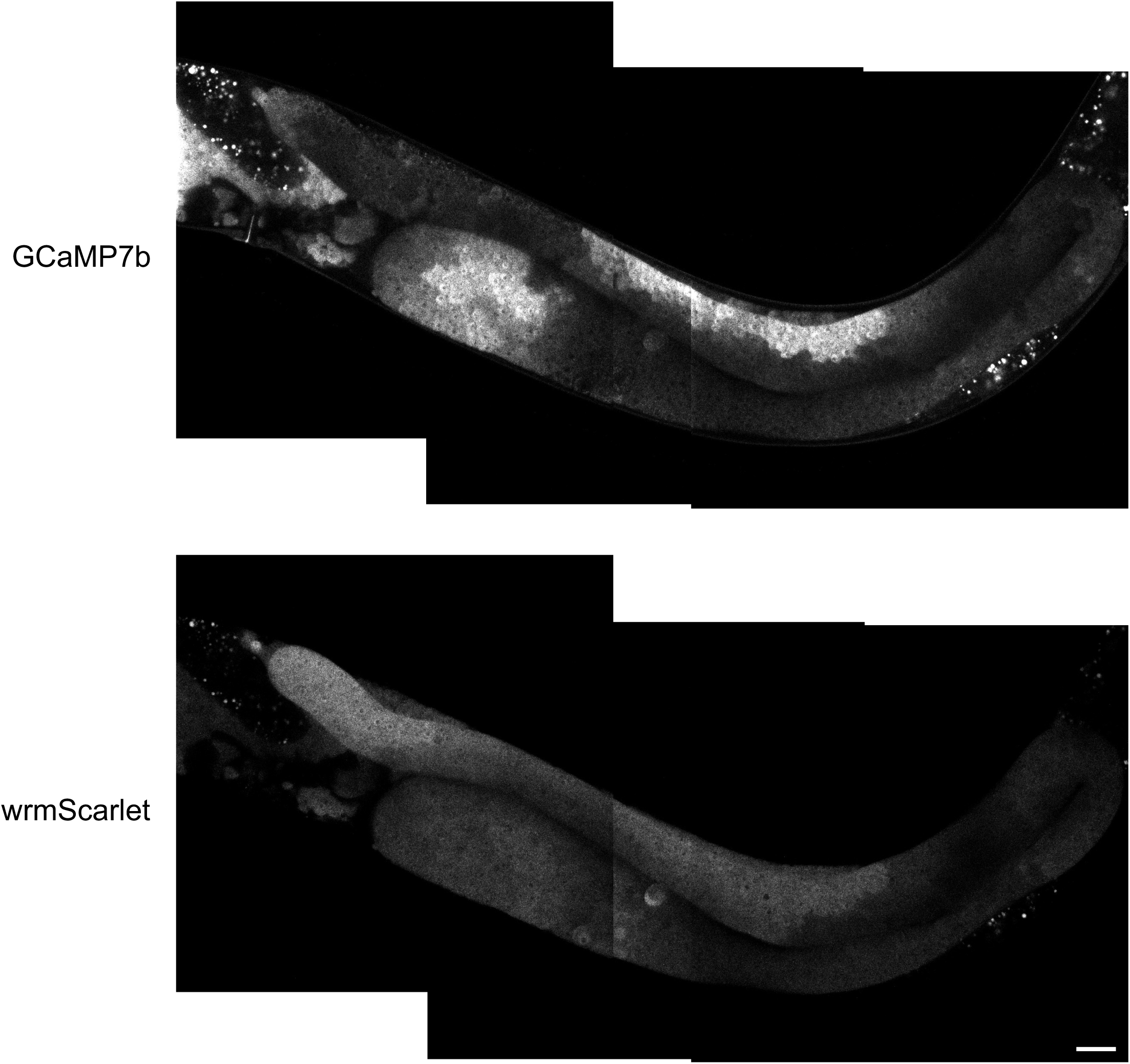
Single channel grayscale images of GCaMP7b and wrmScarlet for the overlaid compound image in Figure 4A. The focal plane varies slightly between the three individual images of the compound image. Scale bar, 20 µm.

## Notes

### Competing Interest Statement

The authors have declared no competing interest.

